# A Neuroscience-Inspired Spiking Neural Network for Auditory Spatial Attention Detection Using Single-Trial EEG

**DOI:** 10.1101/2021.05.25.445653

**Authors:** Faramarz Faghihi, Siqi Cai, Ahmed A.Moustafa

## Abstract

Recently, studies have shown that the alpha band (8-13 Hz) EEG signals enable the decoding of auditory spatial attention. However, deep learning methods typically requires a large amount of training data. Inspired by “sparse coding” in cortical neurons, we propose a spiking neural network model for auditory spatial attention detection. The model is composed of three neural layers, two of them are spiking neurons. We formulate a new learning rule that is based on firing rate of pre-synaptic and post-synaptic neurons in the first layer and the second layer of spiking neurons. The third layer consists of 10 spiking neurons that the pattern of their firing rate after training is used in test phase of the method. The proposed method extracts the patterns of recorded EEG of leftward and rightward attention, independently, and uses them to train network to detect the auditory spatial attention. In addition, a computational approach is presented to find the best single-trial EEG data as training samples of leftward and rightward attention EEG. In this model, the role of using low connectivity rate of the layers and specific range of learning parameters in sparse coding is studied. Importantly, unlike most prior model, our method requires 10% of EEG data as training data and has shown 90% accuracy in average. This study suggests new insights into the role of sparse coding in both biological networks and brain-inspired machine learning.

## I. Introduction

The research of decoding human’s brain activities from Electroencephalography (EEG) signals has progressed by leaps and bounds for various cognitive tasks. Machine learning (ML) algorithms are often used to interpret EEG data. Neuroimaging research has leveraged the advancement in ML algorithms to benefit clicnical practice. These ML methods are based on regression [1], classification [2], or clustering [3] techniques.

In recent years, deep learning has revolutionized the field of machine learning. Deep learning constitutes modern machine learning methods that are based on artificial neural networks [4], [5]. Deep learning methods (DLs) are based on Artificial Neural networks (ANNs) and use non-spiking computing units (named neurons) with non-linear continuous activation functions and set of connection weights (named synapses). Although they are brain-inspired, neural computing and learning rule mechanisms in DLs are different from the brain. In the brain, neurons communicate with downstream neurons by broadcasting trains of spikes. The main challenges to use DL methods are their needs for large amount of training data to avoid overfitting but in many cases is not met [5], and using computationally expensive learning mechanisms, e.g., back-propagation algorithm [6].

Spiking Neural Networks (SNNs) are more biologically models than ANNs. These individual spikes are sparse in time thus information is conveyed by spike timing and spike rates. SNNs are more energy-efficient than ANNs. This feature is appealing for technology, especially for portable devices. Moreover, a SNN encodes and processes the stimulus information through precisely timed spikes trains which makes it a suitable tool for spatio-temporal information [7].

SNNs are tools to simulate neural activities of individual neurons as well as neural population to understand cognitive mechanisms [8]. In addition, SNNs have been used in a number of applications in pattern recognition such as visual processing [9], [10], speech recognition [11], [12] and medical diagnosis [13], [14]. SNNs have been used for the analysis of EEG signals and building Brain-Inspired Brain-Computer Interfaces (BCI) methods [15], for epilepsy and epileptic seizure detection [16], for predictive analysis of various aspects of human behavior during decision making [17], and in the recognition of motor imagery tasks [18]. SNNs usually use bio-inspired learning rules (weight modification) that depends on the relative timing of spikes between connected neurons [19]. Regarding the applicability of SNNs in biomedical data analysis, a SNN architecture called NeuCube has been developed [7] as a general framework for spatio-temporal data modeling that is a fast data processing, and improved accuracy of the EEG data classification.

Pattern recognition in the human brain is done through cortical multi-layer neuronal populations that communicate by spikes. These facts have motivated to develop deep SNNs as brain-like architectures for intelligent machines to implement the tasks that the brain has shown good performance [20], [21]. SNNs have demonstrated good performance in brain signal classification of motor tasks [18] and understanding of spatio-temporal information of EEG during mental tasks [22]. Moreover, these methods may be used in BCI [15] and early diagnosis of degenerative brain diseases [23] and neuro-rehabilitation systems [22].

Theoretical and computational studies suggest that cortical neurons encode sensory information using small number of active neurons with low average firing rate. This phenomenon is named ‘sparse coding’ [24], [25]. Sparse coding confers many advantages including increased memory capacity [26] and efficient pattern separation [27]. In addition, physiological recordings have suggested sparse coding as a ubiquitous strategy in neocortical circuits [28]. The parameters underlying sparse coding of cortical circuits are not fully understood. Unbalanced synaptic excitatory and inhibitory is known to underlie sparse coding in cortical populations [29]. Another parameter is the connectivity in the cortex as an important feature of efficient sensory encoding [29], [30].

Deep sparse coding neural networks have been developed by combining convolutional neural networks (CNNs) and sparse coding techniques for image feature extraction [31], [32] or speech recognition [33]. Classification EEG based on sparse representation and convolution neural network has been proposed to use in BCI applications [34].

Neurophysiology studies have shown that auditory attention can be decoded from EEG data [35]. Recently, ML models have been used to detect the directional focus of auditory attention (left or right direction) solely based on EEG signals [36]. The association of alpha band power of recorded EEG with spatial attention is known [37]. Recently, these findings have been used to detect auditory attention using frequency-domain EEG analysis by CNNs [36]. Moreover, EEG data as time-series have been analyzed using CNN methods [38].

Considering that EEG signals are essentially highly dynamic, non-linear time series data, SNN have by design an advantage over CNN in capturing temporal characteristics of brain activity during different states. In our work, we develop a cortically-inspired classification method to detect directional focus of auditory attention using solely recorded EEG. The developed SNN architecture is based on sparse coding caused by low connectivity of layers as a model parameter and a new learning rule to update the weights during the training phase. The main question that we address in this study is whether it is possible to train an unsupervised SNN-based classification method using small amount of training sample. For this purpose, the dependency of classification accuracy on sparse coding in the neural layers is studied. Moreover, the relationship between spiking properties of neurons and classification accuracy is studied. In the conventional SNN architectures for classification tasks, the firing rate of output neuron(s) is used in the decision process, however, in this method the firing rate pattern of the third layer are used in the test phase to compare the firing rate pattern of EEG samples with rightward trained network and leftward trained network.

The results show the efficacy of this decision making process in EEG analysis. Further studies may show the importance of the decision process in other data types and classification problems.

In our work, synaptic pruning was caused by high values of the learning parameter in combination with low connectivity rate of layers. It induces sparse coding in the spiking neural layer and on another hand, it leads to high classification accuracy. To train the networks, one sample from rightward data and one from leftward data are chosen.

## II. Method

### A. EEG data

In this study, data of a study on 16 subjects listening to audio streams is used [39]. The dataset of the study is publicly available (https://zenodo.org/record/3997352.YJje-WYzb6a). In this dataset, each subject has 24-min EEG recordings for left and 24-min for right. For each subject the data are divided into 10 EEG samples recorded from right ear and 10 EEG samples from left ear. The training as well as test phases use the data of the same subject. In these EEG recording paradigm the competing speakers were located at −90 degrees and +90 degrees along the azimuth direction and there was no background noise. More details on the experiments’ protocol can be found in [39].

### B. EEG preprocessing

The EEG data of each channel were re-referenced to the average response of the mastoid electrodes, then band-pass filtered between 8 and 13 Hz, and subsequently down-sampled to 128 Hz. Finally, EEG data channels were normalized to ensure zero mean and unit variance for each trial. Then the data is normalized into 0 and 1 **(Figure 2a**.)

### C. EEG-based spike encoding

In this study, EEG data are transformed into temporal spike trains using a Threshold-Based Representation method (TBR) [40]. In this method, if the EEG signal exceeds a TBR threshold, a spike occurs. The timing of the spikes corresponds to the time of the change in the EEG data. Different thresholds between 0 and 1 are checked to find the optimal threshold value.

### D. Model architecture and spiking neuron’s model

The EEG data as a time-series transformed into the first neural network composed of 64 neurons (as the number of channels in the EEG data of the study) and spiking train with length of T (ms). The first neural layer is connected to the second layer composed of 100 Integrate and Fire neurons (IF neurons). Different connectivity rate values between these layers are checked to find the optimal value. The second are connected with different value to the third layer that is composed of 10 IF neurons. The connectivity rate value of second and third layer are also considered another model’s parameter. **Table 1**. Shows the parameters value used in simplified IF neuron model in the second and the third layers. **Equation 1**. Shows the IF neuron model.

**Table 1.** Description of IF neuron paramaters used in the SNN model.

|  |  |
| --- | --- |
| $\tau = 0.01$ | the membrane time constant (s) |
| $E_L = -0.065$ | the resting potential (v) |
| $V_{reset} = -0.065$ | the reset voltage |
| $V_{th} = -0.01$ | the threshold voltage |
| $V_0 = -0.06$ | the initial membrane voltage |
| $dt = 0.01$ | the time step (s) |

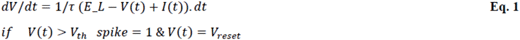

### E. Training phases, learning rule and classification accuracy

To train the network to extract features of the EEG data, one sample from each set of leftward attention EEG and rightward attention EEG are chosen and preprocessed. The training EEG data is presented to the network as the first activated neural network layer that is composed of 64 neurons. Two networks are trained for rightward attention EEG and for leftward attention EEG, independently. The pattern of firing rate of spiking neurons of the third layer for each trained network are measured and saved to use in the decision process **(Figure 1a**.). At the test phase, all EEG samples of right ear and left ear are presented to both right-ear and left-ear trained networks. The firing rate patterns of neurons of the third layer are calculated and compared to the pattern of trained networks. The maximum similarity is used to assign the directional focus of attention to the EEG sample **(Figure 1b**.). For this purpose, Euclidean distance metric was used. The classification accuracy is calculated as the ratio of corrected assigned sample to the total number of samples (here 20 total samples of each subject) **(Figure 1c**.).

**Fig. 1.**
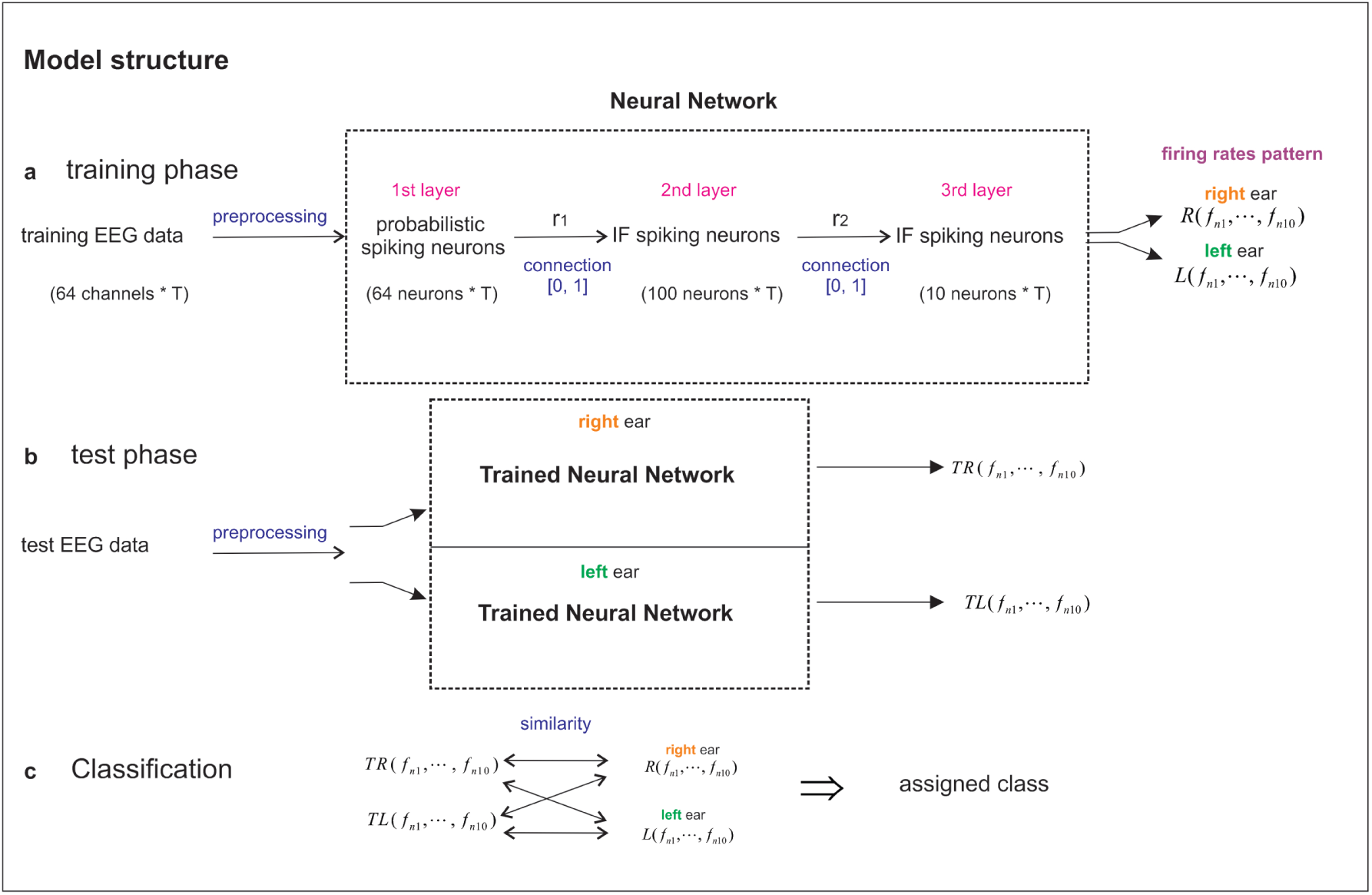
Model structure. **a**. Training phase. EEG samples from left ear and from left ear is preprocessed. EEG data is transformed to 64 spiking neurons as the first layer. The neurons in the first layer are connected to the second layer of 100 IF-neurons with a connectivity value (r1) between 0 and 1. The second layer is connected to the third layer (10 IF-neurons) with a connectivity value (r2) between 0 and 1. Through training, the synaptic weights between the first and the second layer is modified according the proposed learning rule. **b**. In the test phase, the EEG test sample is preprocessed and then is presented into the trained neural networks and the firing rates pattern of the neurons in the third layer is measured. **c**. To assign the directional focus of attention to each test sample, the similarity of firing rates patterns and training samples are evaluated and the maximum value is assigned into the attentional direction.

During the training phase, 1 second of EEG as time window is presented into the second layer. The average firing rate of the neurons in the second layer is measured.

The difference of average firing rate of each pair of neurons in the second layer and the first layer is measured and shown by Δ(*i, j*) **(Equation 2)**. The change in the synaptic weight of the pairs of neurons Δ*W* (*i, j*) is calculated as **(Equation 3)**.

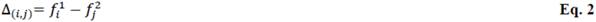

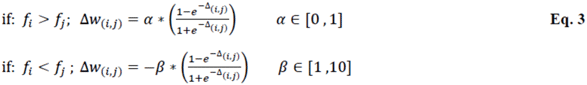

In this learning rule *α, β* are the parameters that are checked to find their optimal values. A threshold is used in order to set synaptic weights lower than the value to zero. Different threshold values are checked to find the optimal value.

### F. Experiments

The classification system to assign class label to an EEG sample of the dataset is checked over different model’s parameters in order to find corresponding values for the optimal classification accuracy. For this purpose, different threshold values of the TBR method are checked. The connectivity values of the layers (r1 and r2) and learning parameters shown in equation 3 are the important parameters that are also of interests from neuroscience point of view **(Figure 1a**.). In addition, we are also interested in studying the possible correlation between sparse coding and classification accuracy. The method is based on single-trial training samples to train the SNN. Therefore, it is highly important to present a method to find the most suitable training samples from rightward and leftward attention EEG before performing calculation of the classification accuracy.

## III. Results

In this section the classification accuracy and the parameters involving in are shown. In addition. For the first time the sparse coding and the neural features of spiking neurons versus classification accuracy is also shown.

### A. Classification accuracy for single training EEG samples

One of the fundamental aims of this study is to use single-trial EEG as EEG samples to train the proposed SNN-based classification method. For this purpose, 10 recorded EEG data from right and 10 samples from left ear data of the subject is used. Different pairs of samples are used in the classification method and the accuracy value is calculated. The optimal threshold of TBR method was calculated as 0.3 and this value is used in all following experiments. Figure 2b. shows the classification accuracy for different possible training sample pairs. The number of x-axis and y-axis are index of samples in the subject’s data. The dependency of the classification accuracy on the pair of selected training samples is clearly recognizable. Maximum accuracy as 90% is obtained for a pair of training samples of a subject. It may be of interests to find the best rightward attention EEG data as well as from leftward attention EEG data. For this purpose, average accuracy over leftward as well as over rightward data are measured **(Figure 2c**.). The maximum average accuracy is assigned to EEGs indexed 13 and 20, for rightward and leftward samples, respectively. In these experiments, the learning parameters *α, β* were set to 8 and 0.1, respectively. In addition, the used connectivity rates r1 and r2 are set to 0.2 and 0.5, respectively.

### B. Dependency of classification accuracy on model’s parameter

Two structural model’s parameters of the developed SNN are the connectivity of the first and the second layer (r1) and the connectivity of the second and the third layer (r2). These parameters are independent and to study the impact of these parameters on the classification accuracy, in each experiment one of them is set while others are considered as variables. First, the connectivity values are set to 0.2 and 0.5 for r1 and r2, respectively. Hence, different learning parameters *α, β* are used and the classification accuracy is measured. **Figure 3a**.shows the classification accuracy on a subject for different *α, β* values. The results demonstrate the important role of high *β* value and low *α* value to obtain high classification accuracy. To study the impact of the connectivity of layers on the classification accuracy *α, β* are set to 0.1 and 8, respectively. The classification accuracy of the developed SNN for different connectivity values between 0 and 1 are shown in **Figure 3b**.. The results demonstrate the important role of low connectivity values (r1 and r2) in obtaining high classification accuracy of the proposed SNN. The importance of these observation in the proposed machine learning method and also in modeling of cortical neural population are presented in the discussion.

**Fig. 2.**
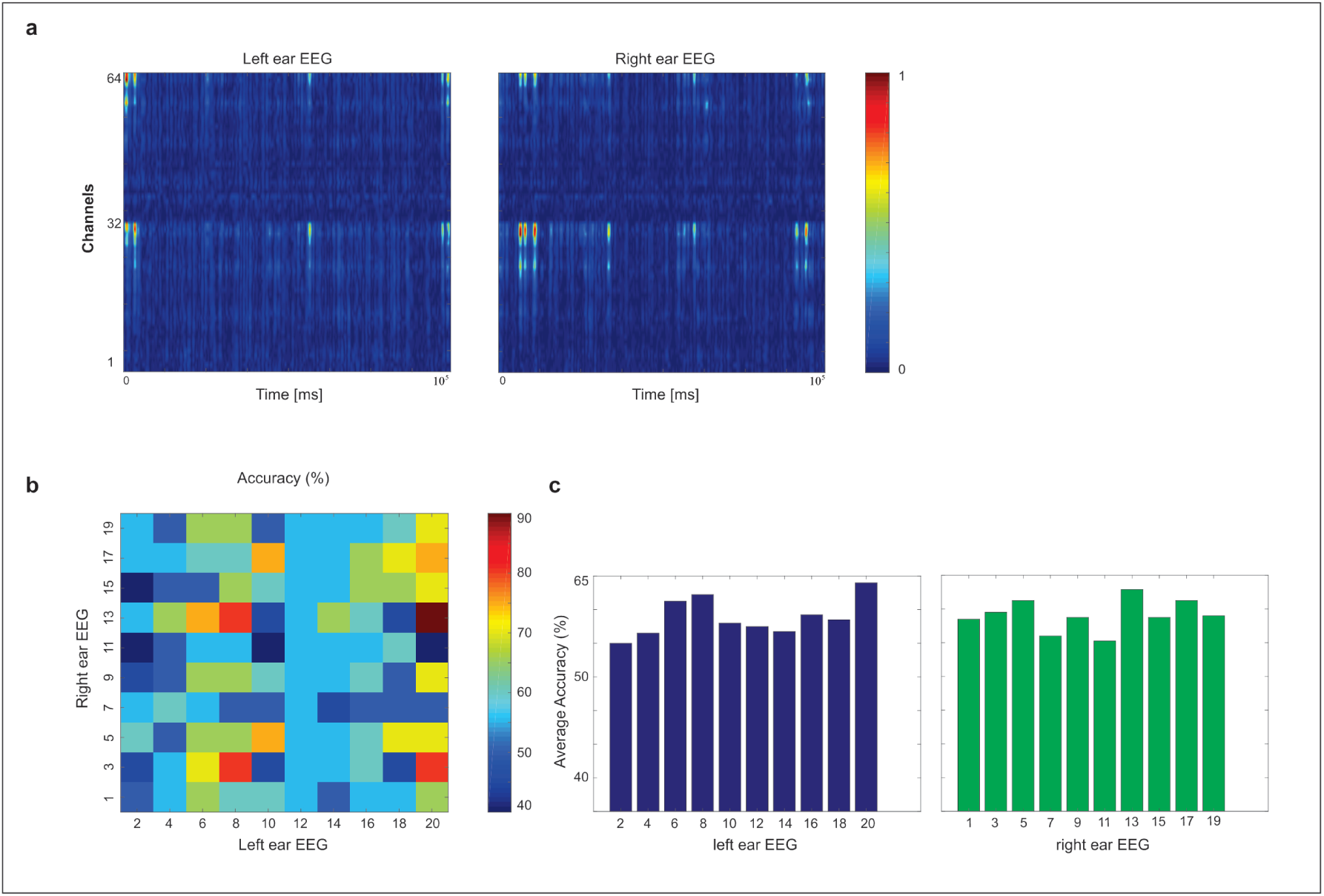
**a**. Two samples of preprocessed and normalized EEG data. **b**. Matrix of classification accuracy for different training samples from rightward and leftward shown by sample index. The accuracy depends on selected training samples. The used connectivity rates r1 and r2 are set to 0.2 and 0.5, respectively. c. To evaluate the best pair of training samples, average accuracy is measured over right as well as left ears data. The best selected samples are 13 and 20 indexes for right and left ear, respectively.

**Fig. 3.**
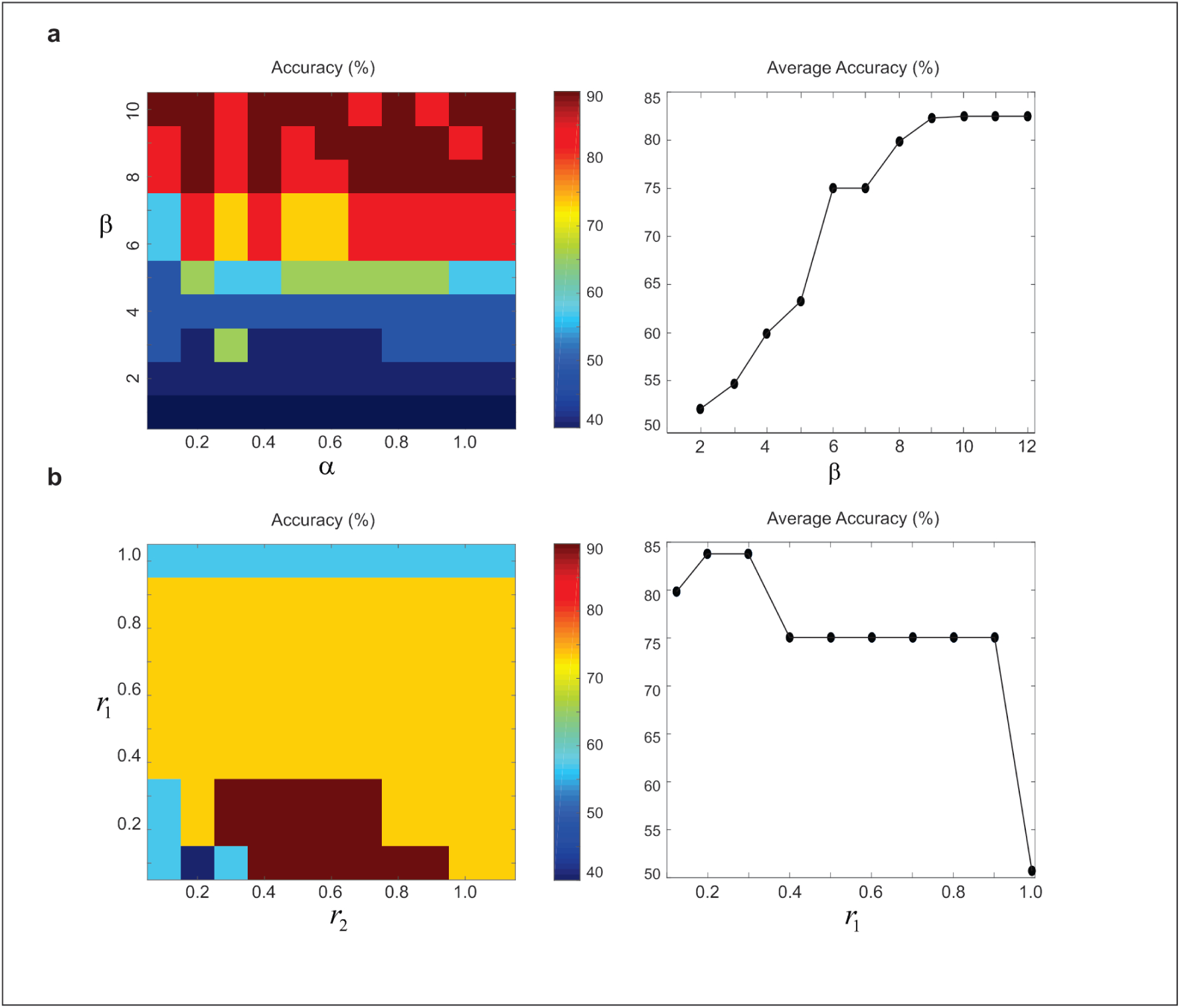
The classification accuracy matrix for different parameters of the model for EEG index 13 and 20. **a. Left panel**. Dependency of the classification accuracy on learning parameters (*α, β*, as increasing and decreasing parameters, respectively). Right panel shows the average accuracy over *α* values. The results the impact of high *β* values on the accuracy. **b. Left panel**. The classification accuracy matrix for different connectivity values of the model. Right panel shows the average accuracy over the connectivity of the second and the third layers. The result shows the impact of sparse coding (low connectivity values) in the second layer on the classification accuracy of the model.

### C. Synaptic weights and sparse coding changes through training

Through presenting EEG training samples to the network, connectivity values r1 and r2 are set to 0.1 and 0.5, respectively. The synaptic weights of the connected neurons of the first layer and the second layer are initially selected randomly as values between zero and one and through the training of the network are changed according to the learning parameters. *α* is set to 0.2 and the synaptic weights for different *β* values are measured. Increase in *β* value results in more deleted synapses at the end of training the proposed spiking neural network **(Figure 4a)**. The changes in the synaptic weights between the first layer and the second layer leads to change in spiking rates and number of activated neurons of the second layer. Consequently, such changes in spiking rate and the number of active neurons in the second layer leads to the change in the firing rate of neurons in the third layer. Using higher *β* values result in sparse coding in the second layer and the third layer **(Figure 4c)**.

**Fig. 4.**
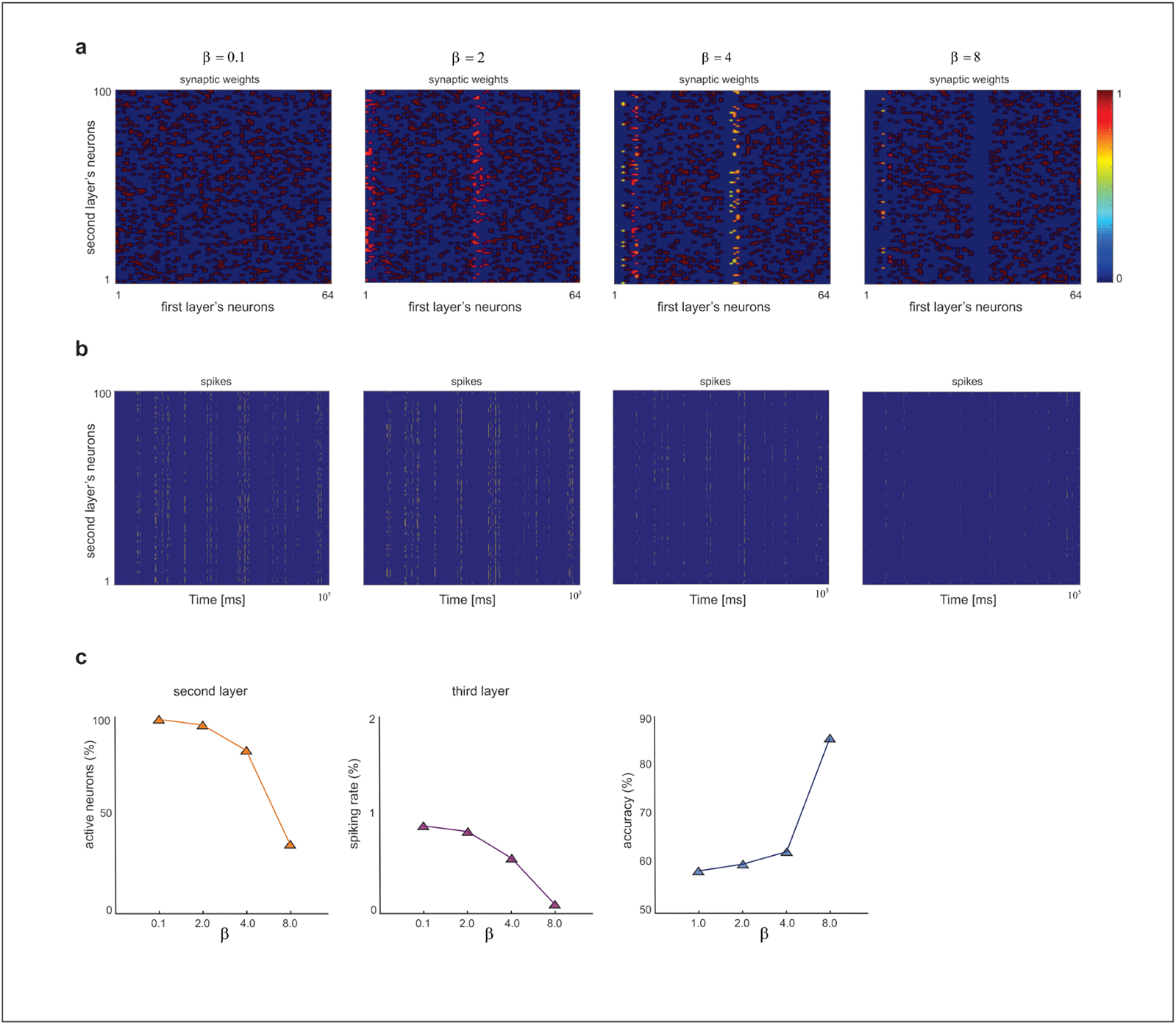
**a**. The dependency of the synaptic weights modification on *β* parameters value for *α* = 0.1. Higher *β* values leads to more deleted synapses through training phase. **b**. The impact of *β* parameters value on spiking rates and sparse coding of the model. **c**. Left panel shows the impact of *β* values on the number of activated neurons of the second layer. Increase in *β* value leads to decrease of activated neurons. Middle panel. In addition, increase in *β* value results in sparse spiking in the neurons of the third layer. Right panel. The average classification accuracy for different *β* values over *α* values.

**Figure 4c** also shows relation between sparse coding in the second layer and third layer with classification accuracy using pair of rightward and leftward attention EEG samples corresponds to the maximum obtain accuracy.

### D. Spikes distance of the neurons for different learning parameter values

Using high *β* values induce change in spiking activity of neurons in the second and third layers while it leads to increase in classification accuracy. We are interested to study dependency of the classification accuracy and the distance of spike trains of neurons (or dissimilarity of spike trains) as a measure of neural synchrony. **Figure 5a, 5b** shows the distance of spike trains of neurons of the second and the third layers for incremental *β* values. Higher *β* values induce increase in distance of spike trains in both neural layers. **Figure 5c** shows the average distance of spikes trains over total neural population in the second and the third layer and compare them with the classification accuracy for different *β* values. It demonstrates that the increase in classification accuracy is associated with decrease in average distance of spike trains in both second and third neural layers. In all these experiments *α* is set to 0.2.

**Fig. 5.**
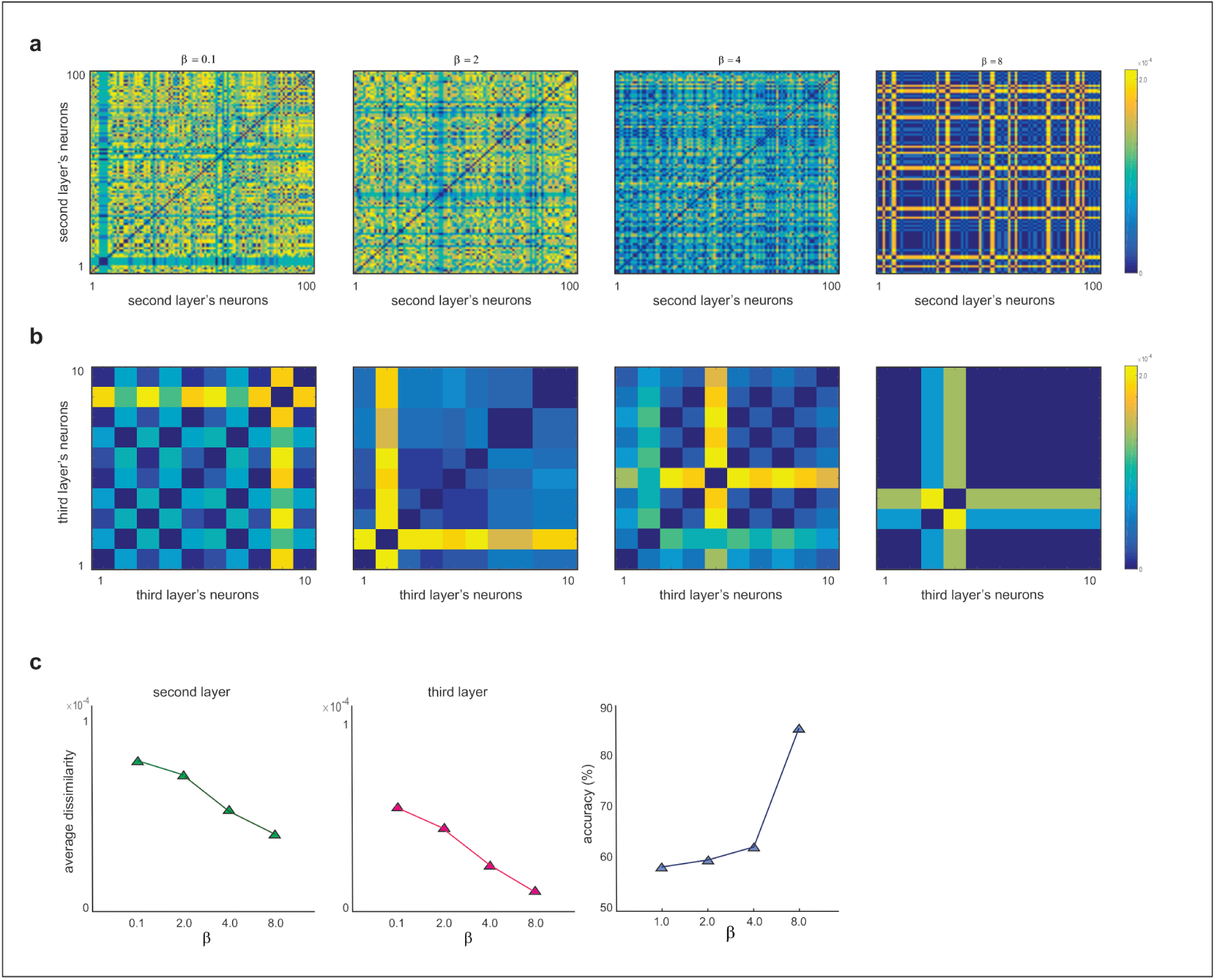
**a**. Dissimilarity of spiking pattern of neurons in the second layer and in the third layers (**b**.) for different *β* values. **c. Left and middle panels**. Average dissimilarity of the second and third layers for different *β* values. Increase in *β* values leads to decrease in average dissimilarity of spiking neurons in both second and third layers. **Right panel**. Average classification accuracy for different *β* values (*α* = 0.1). Increase in *β* value results in raised accuracy while decreasing in dissimilarity of spiking patterns of the neurons.

### E. Predicting the best EEG samples to train the SNN

In this study, all possible pairs of training samples (one from leftward attention EEG data and another from rightward attention EEG data) from a subject were considered. The classification accuracy were calculated by using each pair. Assigning an EEG sample to rightward or leftward classes is based on measuring and comparing the similarity between firing rate of neurons of the third layers of the test sample with trained network. Therefore, it is expected that if the network is trained using each samples from rightward or leftward, independently, the best training samples have third layer’s firing patterns with low average dissimilarity with other firing patterns in each class. To test this hypothesis, the information shown in **Figure 2b** is plotted versus the proposed measured dissimilarity for rightward EEG and leftward EEG, independently. The results show negative correlation between these two measures for both rightward and leftward of a subject. In addition, the best training samples from rightward and leftward EEG data are shown by red stars (**Figure 6** left and middle panels). The average correlation over all subjects are −0.45 and −0.51 for rightward and leftward EEG data, respectively (**Figure 6** right panel). These results propose a method to select a subset of the data as the best candidates to use in training process.

**Fig. 6.**
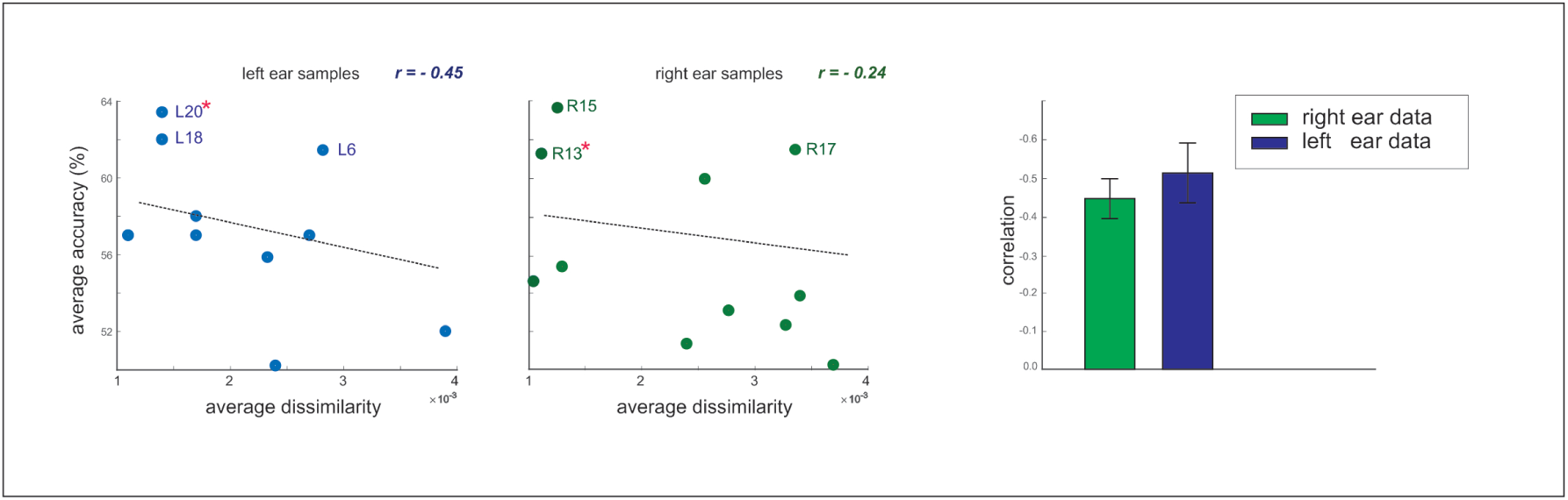
Predicting the best candidates for training samples. For rightward and leftward EEG samples each sample is used in the process of training with predefined models’ parameters. The similarity of third layer ‘firing rates patterns’ are calculated and are drawn versus average classification accuracy shown in figure 2c. The results show negative correlation with −0.24 and −0.45 values for right ear and left ear EEG, respectively. The best candidate are shown with their labels. The samples corresponding to maximum accuracy shown in figure 2b are shown with red stars.

### F. Classification accuracy over different decision window lengths

In order to apply the classification methods for online auditory attention detection, short decision window to make decision on the test sample is expected. Therefore, the classification accuracy of the proposed SNN-based method is measured for different length of decision window. **Figure 7** shows the results on the decision windows between 20 seconds and 1 second. A decrease in the decision window leads to decrease in the average accuracy. Therefore, the proposed classification method needs long decision window to demonstrate high classification accuracy.

**Fig. 7.**
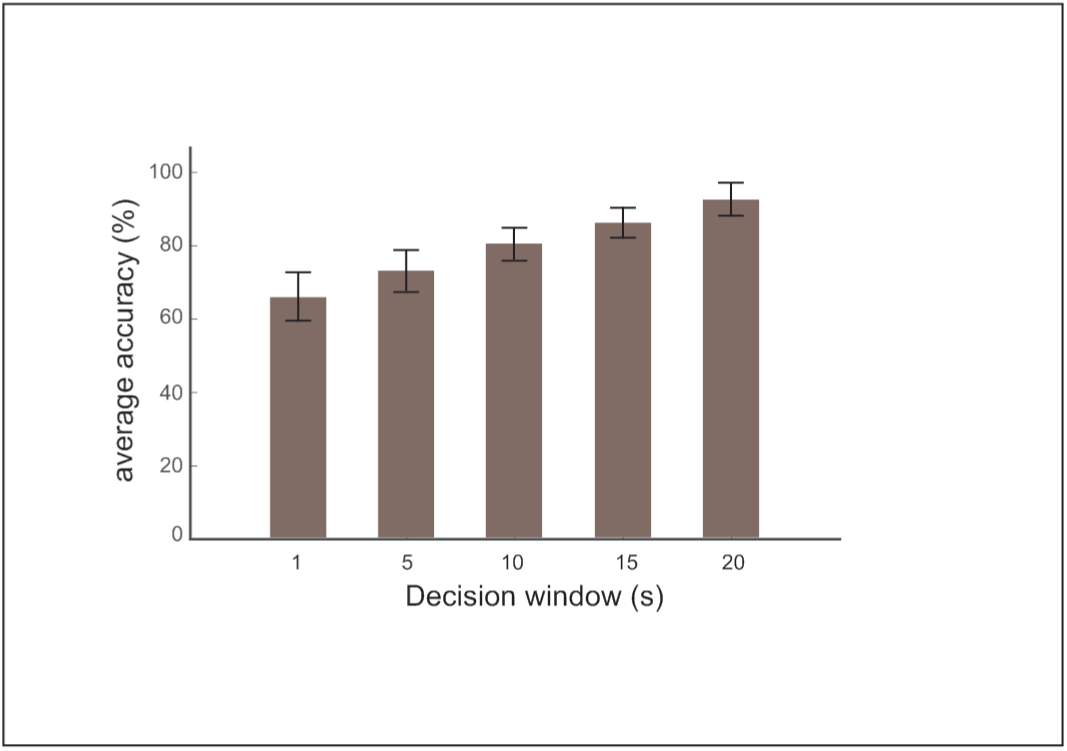
Classification accuracy of the proposed SNN model over decision windows from 1 to 20 seconds EEG data. Shorter windows results in decrease in accuracy over the subjects.

## IV. Discussion

Inspired by the findings that directional focus of the auditory attention is encoded by cortical neurons, a cortically-inspired model was developed to detect the attention activities manifested in brain signals. In a research paradigm of auditory attention decoding, audio stimuli is presented to rightward or leftward of the subject which is simultaneously recorded from scalp electrodes from many brain regions. Therefore, recorded EEG from the subject stimulated from rightward and leftward may have very similar pattern. The aim of auditory attention decoding is to extract features of the training EEG data to use in determining the directional focus of attention using test EEG samples to be either rightward and leftward classes. In this study, a framework presented to select training samples prior to classification process **(Figure 6)**. For this purpose, the EEG samples of rightward (10 samples per subject) are used to train the network independently. The firing rate patterns of the third layer obtained by presenting each EEG sample (as training sample for rightward) is measured. The similarity matrix of 10 obtained patterns are calculated and the EEG sample corresponding the maximum similarity to other EEG samples are selected as the best training sample for rightward. The same process finds the best EEG sample for leftward EEG data. The results on different subjects demonstrate 75 % success of the method in finding the best training samples. The results demonstrate the dependency of classification accuracy on the selected pair of training samples **(Figure 2b)**.

Conventional deep learning methods (e.g., CNN) requires large training dataset. Although our method shows lower average classification accuracy (90 %) compared to the study in [36] (that is 94 %), it needs only 10 percent of the data as training sample and hence proposed a novel class of spiking neural networks for applications where small training sample is provided. Our method typically requires less training data, is more robust and stable, and is computationally cheaper. Therefore, the proposed method may be of interests for deep learning researchers where for a specific problem there exist small set of training samples, especially for EEG dataset. Hence, our SNN architecture should be checked in different applications to prove its efficiency as a novel neural network classification method that needs small amount of training data while it tries to find the suitable training samples.

For real applications, it is required to make decision on the direction of attention using EEG data as less as possible. Although in all deep learning methods using short decision windows results in decrease in classification accuracy, our method using short decision windows cause drastically decrease in the classification accuracy. This limits the application of the proposed SNN in neuro-rehabilitation devices. Therefore, future works should be concentrated on overcoming on this challenge.

To our knowledge, SNN architectures have not been developed for auditory attention decoding yet. The proposed SNN instead of many neural layers that are usually used in CNNs, is composed of three layers where two layers are spiking neurons. Therefore, its computational cost is low compared to deep learning methods. In addition, it belongs to unsupervised classification methods, so it does not need back-propagation based learning mechanisms. This learning rule is inspired by back-propagation of spikes from soma to the dendrites in response of biological neurons to external stimuli and modifies the synaptic weight between pre-synaptic and post-synaptic neurons [41], [42]. In the proposed SNN that has two spiking neural layers, features of training EEG data is extracted as changes in the synaptic weights of connection between the first and second neural layers through training process. The *α,β* parameters of the proposed learning rule determines the increase or decrease in the synaptic weights. Using high *β* values lead to deletion of weak synapses and hence helps to extract important features of the EEG. This process is inspired by cortical neurophysiology and is called ‘synaptic pruning’. Synaptic pruning is commonly observed during the development of human brain [43]. Regarding the important role of synaptic pruning in neural development, some spiking neural networks have tried to use it in their architecture. One example of this class of SNNs is a model for neural coding that is inspired by sparse coding and random connectivity of by real neural circuits where structural changes in the random connectivity induced by pruning process. In this work [44] random sparse connectivity has been presented as a key principle of cortical computation. In order to implement pruning, a STDP (Spike-Time Dependent Plasticity) model has been proposed [45]. In our work, spike correlation between pre-synaptic and post-synaptic neurons are used such that synapses with low correlation or uncorrelated spiking activity are pruned. In our work, synaptic pruning was caused by high values of *β* as learning parameter in combination with low connectivity rate of layers. It induces sparse coding in the spiking neural layer and on another hand, it leads to high classification accuracy. To train the networks, one sample from rightward data and one from leftward data are chosen.

In the conventional SNN architectures for classification tasks, the firing rate of output neuron(s) is used in the decision process, however, in this method the firing rate pattern of the third layer are used in the test phase to compare the firing rate pattern of EEG samples with rightward trained network and left-ear trained network. The results show the efficacy of this decision making process in EEG analysis. Further studies may show the importance of the decision process in other data types and classification problems.

Some deep learning methods have been proposed for auditory attention decoding [38]. Recently, a deep learning method has been proposed that takes single-trial EEG data and the spectrogram of the multiple speaker as input and classifies the attention to one of the speakers. This deep learning architecture is a joint CNN with LSTM model (Long Short Term Memory) [46]. The method has demonstrated 77.2% decoding accuracy at trials with decision window of three seconds (the amount of EEG sample used to make a decision). Recently, a CNN architecture was used as a classifier and CSP was used for EEG signal enhancement under different auditory attention paradigms. This method has demonstrated 94% classification accuracy [36]. Common Spatial Pattern (CSP) is a method to determine auditory attention direction solely based on recorded EEG. Riemannian geometry-based classification method has been developed as an alternative for CSP approach. In this approach, covariance matrix of EEG data is directly classified with taking its Riemannian structure into account [47]. Computational, and experimental studies suggest that sensory information is encoded by small number of active neurons of a neuronal population at any given time. Computational studies have shown the role of sparse coding in memory storage in hippocampus [27] or cortex [48]. Experiments have shown that different connectivity pattern of different cortical cells is essential to encode sensory information from different modalities [49], [50] and to understand many neuronal responses [51]. Our results also suggests the dependency of the classification accuracy on low connectivity of layers that is also associated with sparse coding in the developed spiking neural layers **(Figure 3)**.

Our model that has been inspired by these observations in cortical neurons, interestingly illustrates a relation between the classification accuracy and the similarity of the neural spike trains of the second and third neuronal layers that is also depends on sparse spiking of neurons. Such similarity of the neural spikes can be considered as a measure of neural synchrony. The simulations of this study assign important role for sparse coding in both brain-inspired machine learning studies and information encoding in cortical neurons where sparse coding is a known neuronal population feature. In summary, the developed SNN-based classification method can be trained with single trial EEG data per rightward and leftward directions while it has shown 90% classification accuracy. Low amount of required training data for neural network-based classification is highly important for applications in complicated data analysis where small amount of training data is accessible. To our knowledge, it is the first time sparse coding SNN is used in classification purposes. In this SNN a firing rate based learning rule and synaptic pruning were used while the role of different connectivity rate of layers in sparse coding and its correlation with classification accuracy was studied. We think that the SNN presented in this study offers new insight into both the role of sparse coding in brain-inspired machine learning and possible mechanisms of information processing in cortical neurons.

## Acknowledgment

This research work is supported by Programmatic Grant No. A18A2b0046 and A1687b0033 from the Singapore Government’s Research, Innovation and Enterprise 2020 plan (Advanced Manufacturing and Engineering domain).

The work by Faramarz Faghihi is funded by the Deutsche Forschungsgemeinschaft (DFG, German Research Foundation) under Germany’s Excellence Strategy (University Allowance, EXC 2077, University of Bremen, Germany).

